# The effect of climate change on the distribution of Canidae

**DOI:** 10.1101/2021.07.19.452957

**Authors:** Lucas M. V. Porto, David Bennett, Renan Maestri, Rampal S. Etienne

## Abstract

Land use by humans and climate change have been seriously affecting the distribution of species resulting in a quarter of all known mammals currently threatened with extinction. Here, we modeled the present and future potential distributions of all 36 extant Canidae species to evaluate their response to future climate scenarios. In addition, we tested if canids were likely to experience evolutionary rescue, which could allow some species to adapt to climate change. Our results suggest that global warming will cause most species to lose or maintain their ranges, while a few will have the potential to benefit from future conditions and considerably expand their geographic distributions. Some canids have the potential to experience evolutionary rescue, but *Atelocynus microtis* and *Chrysocyon brachyurus* are two concerning cases that do not show this capacity to adapt given the current pace of climate change. We also reveal that most Canidae hotspot regions are outside protected areas, which may be useful for the identification of key areas for conservation.

## 1 INTRODUCTION

The pace of climate change induced by humans is much faster than predicted previously (Pimm et al., 2014). Ceballos et al. (2015) showed that the rate of vertebrate species loss over the last century is up to 100 times higher than the background extinction rate. This anthropogenic pressure causes habitat loss and increased competition from invasive organisms (Butchart et al., 2010), which leads to species extinction (Ceballos et al., 2015; May & Lawton, 1995), and thus has serious impacts on global biodiversity. Over the last decades, several studies have shown how human impacts affect the structure of ecosystems and how these changes can backfire and affect humans negatively with floods, fires, air pollution, heat waves, and vector-borne diseases (Bellard et al., 2012; Cardinale et al., 2012; Goberna et al., 2014; Kortsch et al., 2015; Nadeau et al., 2017; Parmesan & Yohe, 2003; Pecl et al., 2017; Woodward et al., 2010). Some species are more susceptible to extinction than others due to their traits, including: reproductive rate, habitat specialization, body size, and geographic range (Davidson et al., 2009; Fritz & Purvis, 2010). Therefore, understanding how species are going to respond in future scenarios of climate change is necessary to predict the impact of the loss of certain species on ecosystems, and it will be useful for conservation of biodiversity.

Until recently, evolution was thought to play no substantial role in a population’s resilience when facing a rapid environmental change (Ferrière et al., 2004). The common idea was that a population in decline, exposed to a deteriorating environment, would become extinct. However, Gonzalez et al. (2012) and Bell (2013) coined and matured the idea of evolutionary rescue (ER). In an ER scenario, adaptive processes could be triggered in some resistant individuals of the population under environmental stress, allowing them to rapidly proliferate and counter the rate of decline of the population, thereby changing our perspective on communities with populations that are threatened with extinction (van Eldijk et al., 2020).

The most used tools to evaluate how species are dealing with climate change are ecological niche models (ENMs) (Araújo & New, 2007; Ehrlén & Morris, 2015; Elith et al., 2010; Guisan & Thuiller, 2005). ENMs use mathematical modelling of a species’ relationship with environmental variables, and predict habitat suitability for that species based on known occurrence data (Araújo et al., 2011; Guisan & Thuiller, 2005). ENMs based on climate data have proven extremely useful in assessing the effectiveness of the distribution of protected areas (Catullo et al., 2008), assessing species vulnerability to local land-use changes (Santos et al., 2013), predicting distributions of rare species (Marino et al., 2011; Rheingantz et al., 2014), and predicting possible responses to climate change by species and ecosystems (Moor et al., 2015; Sobral-Souza et al., 2018).

However, the use of ENMs with climatic variables alone has been debated in several studies (Diniz-Filho et al., 2019; Elith et al., 2010; Synes & Osborne, 2011), mainly because ENMs do not incorporate intrinsic characteristics of the populations, relying on the idea that all the mechanisms that affect species` distributions are captured by the environmental data (Diniz-Filho et al., 2019). However, niche models that use traits (morphological and physiological) or genetic data are complex and do not work well when the niches of several species are modeled simultaneously (Norberg et al., 2012).

The attempt to predict responses of species to climate change is further limited by uncertainties surrounding climatic predictions - with slight differences existing between different general circulation models (GCM) - and by uncertainties about the possibilities of measuring evolutionary rescue.

Recently, Diniz-Filho et al. (2019) applied a macroecological framework to estimate responses to evolutionary change and the likelihood of evolutionary rescue; they proposed the *H* value (Haldanes) to estimate the evolutionary change required by species to maintain their populations in future environmental scenarios, giving a biological and evolutionary meaning to temperature variations that species will experience. According to the framework proposed by Diniz-Filho et al. (2019), the greater the variation in temperature between present and future, the greater the *H* value, and consequently, the more difficult it is for the species to experience an ER scenario. Likewise, the fewer generations the species can have until the future, the higher the *H*. In short, the smaller the temperature difference and the larger the number of generations, the more likely it is for evolutionary rescue to happen, and for a species to persist in the face of climate change.

WWF (2018) showed an average 60% decrease in vertebrate populations, and a quarter of all known mammals are currently threatened with extinction (IUCN, 2020). Within this group, the canids (family Canidae) is an excellent group to test the impacts of climate changes on future distributions, as they are distributed in all continents, except Antarctica (Sillero-Zubiri et al., 2004; Wang & Tedford, 2008), and because as medium-large mammals they are more prone to extinction than smaller species (Rija et al., 2020). Like other species, canids are affected by the consequences of urbanization and climate change: coyotes (*Canis latrans*) and red foxes (*Vulpes Vulpes*) have been observed in urban areas in North-America (Lombardi et al., 2017; Mueller et al., 2018; Poessel et al., 2013, 2017), while the red fox has invaded a habitat in northern Europe that was previously occupied only by the arctic fox (*Vulpes lagopus*) (WWF, 2018). Understanding how canids are affected by changes in the landscape, and being able to predict their future distributions is essential to outline conservation strategies for different species.

Here we use climate-based ENMs to: 1) model the distribution of all canids under present climate conditions, 2) predict possible changes in Canidae distribution under climate change in the next 54 years (2075), and identify species at risk of losing some or all of their current range, but also assess if some species could enlarge it; and 3) identify which species are most likely to adapt to changing climatic conditions and therefore avoid the negative effects of temperature change.

## 2 MATERIALS AND METHODS

### 2.1 Occurrence and environmental data

Species occurrence data for all canids were taken from VertNet (Constable et al., 2010) and the Global Biodiversity Information Facility (GBIF, 2020) online databases. The number of occurrence points is shown in Tables S1, and cover all the known distribution of the 36 species used here (which correspond to 100% of the living Canidae species). We spatially filtered the data using SDMToolbox 2.0 (Brown et al., 2017), in ArcGis 10.3.1 (Environmental Systems Resource Institute, 2019), to remove duplicate occurrence points. As there are different classifications for the Canidae family in relation to the number of species (Bardeleben et al., 2005; Perini et al., 2010; Zrzavy & Ricánková, 2004), here we use the most recent canid phylogeny proposed by Porto et al. (2019) to define which species of Canidae (*n* = 36 - Table S1) would be considered here to model their potential distributions.

For environmental variables, we downloaded a digital elevation model (DEM) (IUCN, 2019) and the standard 19 Worldclim bioclimatic variables for the present and future (2075) (Hijmans et al., 2005). In addition, we used the distance to freshwater as a variable, which we measured using the Natural Earth River and lake maps, and the Euclidean distance tool in ArcGis. To clarify the environmental data we masked the variables and imported them into R 4.0-2 (R Development Core Team, 2020) and tested for multicollinearity using variance inflation factor (VIF) tests with the package regclass 1.6 (Petrie, 2020) and pairwise plots. Highly correlated variables (VIF score > 10 or Pearson correlation > 0.7 respectively) were eliminated one at a time, starting with the variable(s) deemed to possess the least ecological relevance based on the VIF tests.

### 2.2 Ecological niche modelling

ENMs for the present were performed using the R package SDM 1.0-89 (Naimi & Araújo, 2016). To model species’ niches for the present, we generated 10.000 random background points within a mask equivalent to the species’ known IUCN ranges, buffered to 220 km (or approximately two decimal degrees), producing a presence-absence matrix of species within defined grids cells (pixels). We built ensembles (objects with a weighted averaging over all predictions from several fitted models) of four different models: Maxent, Support Vector Machine (SVM), Random Forest (RF), and Boosted Regression Tree (BRT). For all models, we used 90% as training data and 10% were retained as test points. Models were only accepted if they had acceptable True Skill Statistic (TSS - calculated as the sum of specificity + sensitivity – 1) and Area Under the Curve (AUC) values (0.7 being the minimum accepted AUC, 0.6 the minimum TSS (Allouche et al., 2006)). We used both TSS and AUC to evaluate the models because they assign different weights depending on the sample size of the data used (Guisan et al., 2017), and hence we believe our results to be more robust if both criteria are met.

In order to verify whether the ENMs and IUCN polygons agree, we compared the current distribution maps of all species of canids available at IUCN against the maps created here through ENMs. IUCN maps were generated by minimum convex polygons, which represent the realized niche of the species, while the ENMs here bring a more detailed notion of their fundamental niche.

We modelled the future distribution of species based on the most pessimistic climate scenario for the year 2075 (RCP 8.5 - Representative Concentration Pathway) from IPCC (2007). We chose this scenario because it seems to have become the most realistic one over the last years, and can even be under-estimating future concentrations of atmospheric carbon (Christensen et al. (2018). RCP 8.5 assumes high global CO_2_ concentration, a high rate of human population growth, and an increased use of energy and land. We used an ensemble of three General Circulation Models (GCMs): Access1.0 (exhibits a high skill score with regard to historical climate), HadGEM2 (has a good representation of extreme El Niño events), and MIROC5 (also has a good representation of extreme El Niño events, and represents all RCPs scenarios well). Maps of suitability (present/future) are shown on a continuous scale to better visualize the potential distribution of species.

### 2.3 Evolutionary rescue calculations

*H* values were calculated for each of the 36 canids to predict whether they can adapt to climate change and prevent the loss of their habitat. We assumed that temperature is a representation of the species’ niche (tolerance) most closely reflecting climate change. For each species, changes in maximum temperature of the warmest month (Bio05) across the entire range were estimated, and the temperature change in each cell was calculated as the average of the future temperature (in the warmest month) minus the average of the present temperature (in the warmest month). Following Diniz-Filho et al. (2019), *H* values were calculated using:

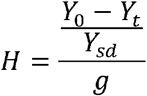

where *Y*_0_ is the mean temperature at the present, *Y_t_* is the mean temperature at time *t* in the future, *Y_sd_* is the standard deviation of the present temperature tolerance (assuming a constant variance between generations), and *g* is the number of generations between present and future. The generation lengths for all canid species was compiled from the Animal Diversity Web (ADW - Myers et al., 2018) and PanTHERIA (Jones et al., 2009) (Table S1). The higher the value of *H*, the greater the rate of evolutionary change needed for a species to experience ER, and consequently, the more difficult it is to maintain its population facing a climatic change scenario.

For the evolutionary rescue analyses, we used the threshold maps (binary) for each species, produced with the suitability maps because they show presence/absence values based on the specificity and sensitivity of the model (Liu et al., 2015).

## 3 RESULTS

### 3.1 Ecological niche modelling

All ENMs produced acceptable accuracy values for TSS and AUC. After testing highly correlated variables, only five were not excluded and were used to model canid niches, they are: distance to freshwater (DIST), the maximum temperature in the warmest month (Bio05), precipitation in the driest month (Bio14), elevation (DEM).

To check the reliability of the ENMs we compared their predictions on the present distributions with the actual current distributions according to IUCN polygons (realized niche) (Appendix - Figure S1 – S36). With the exception of a single species, *Canis lupus*, the distribution polygons fall within the areas that the ENMs demonstrate to be suitable for the species to occupy (fundamental niche). Species richness maps for the present generated by ENMs (Figure 1) and by polygons (Figure S37) show very similar patterns of species overlap, generally maintaining the same hotspot locations in the Middle East + Northeast Africa region and western part of the USA. However, there is an exception: the richness map based on polygons shows the presence of Canis lupus in the Middle East region towards India, but this is not predicted by the ENMs (see Discussion). Because of the high similarity our ENM predictions seem highly reliable, and we therefore compare our future ENM predictions with ENM predictions for the present, as they are better comparable (both describe the fundamental niche).

**FIGURE 1.**
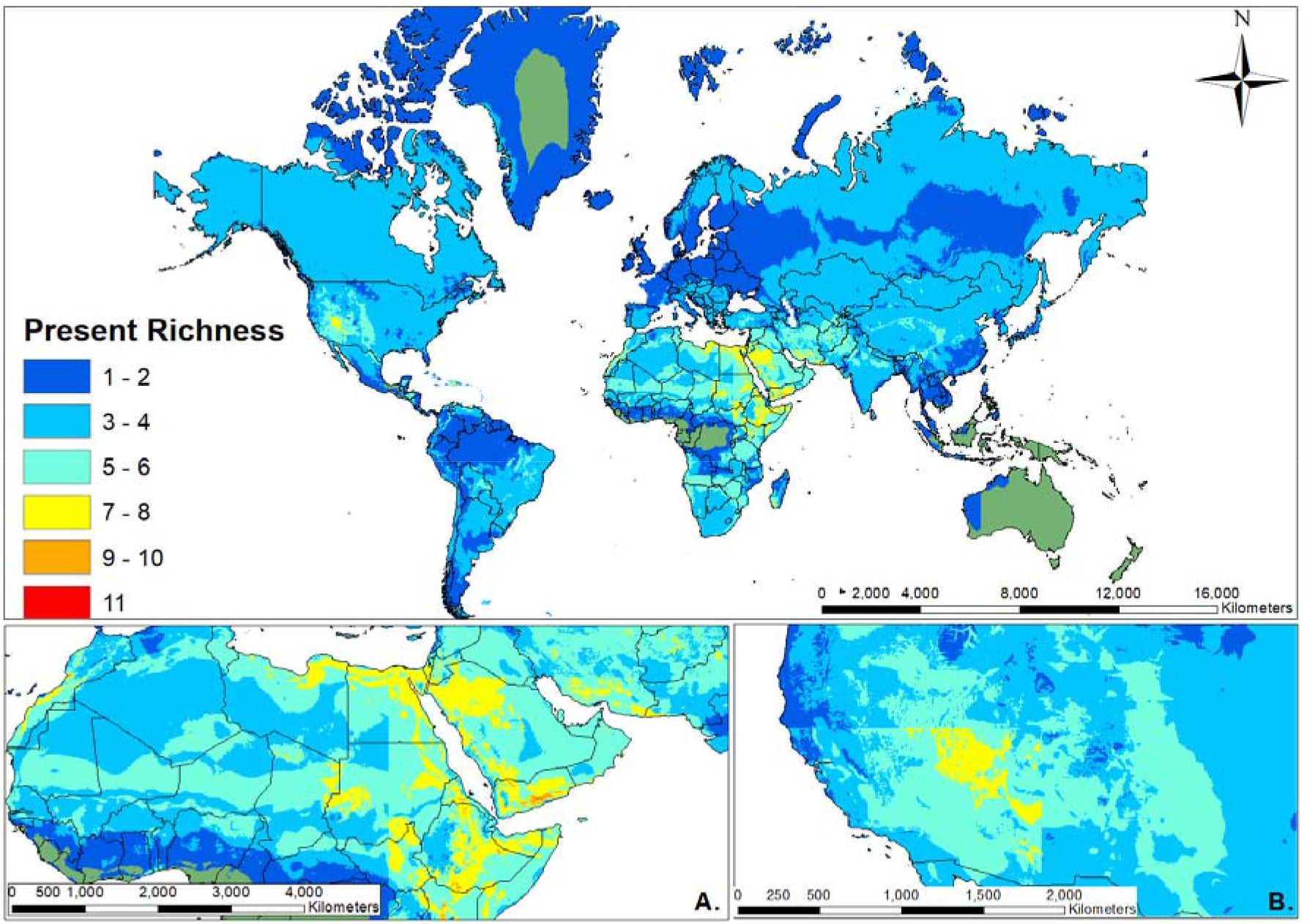
Species richness map of Canidae for the present produced by ENM. The richest areas (hotspots) were identified in the Middle East + Northeast Africa region (A) and western part of the USA (B). The legend on the left shows the number of overlapping species.

Our models indicated that 27 species were predicted to experience range contractions under climate change, while 9 were predicted to expand in range overall (Table S2). In all three Canidae clades (wolves, foxes, and South-American canids), we find that most species will contract their ranges, and a few will expand their ranges (Figure S38A–S38C). We discuss them now in more detail.

The South-American canids (Figure S38C), *Atelocynus microtis*, *Lycalopex fulvipes*, and *Lycalopex sechurae* are predicted to see future climate suitability fall below their modelled threshold across their entire ranges (Table S2), losing a large part of their geographical distributions (Appendix - Figures S39, S40, and S41). In contrast, *Cerdocyon thous* is the only South-American canid that was predicted to have a considerable expansion in its geographical area under future conditions; moreover, the ENM predicts that *C. thous* will occupy areas within the Amazon Forest not inhabited before (Table S2, and also see Appendix - Figure S38C and Figure S42).

In the clade of wolves, *Canis latrans* and *Canis rufus* are probably going to lose a large part of their ranges in North America, while *Canis anthus* and *Canis lupus* are expected to increase their distributions, mainly in desert areas such as the Middle East, for both species, and the deserts in the USA for *C. lupus* (Table S2, and also see Appendix - Figure S38A, Figure S43, and Figure S44).

Some of the fox species are predicted to suffer severe losses in their ranges (Table S2 and Figure S38B). Among them, *Urocyon littoralis* stands out: even though it is considered an endangered species at the moment, the ENMs predicted that *U. littoralis* will lose 28.6% of its (small) current distribution (Table S2). *Vulpes chama*, *Vulpes bengalensis*, and *Vulpes velox* also were predicted to have a considerable decrease in their geographical ranges. By contrast, *Vulpes corsac*, *Vulpes vulpes, Otocyon megalotis*, and *Urocyon cinereoargenteus* will probably experience range expansions under future climatic conditions. In fact, the ENM predicted that *V. vulpes* will increase 5.7% of its distribution, inhabiting new areas such as the Middle East, Northern Canada, and Greenland (Appendix - Figure S45).

The richness map of the present (Figure 1) shows that the overlap of different species is very low around the planet. The map also points to two hotspot areas for canid diversity, one in the western part of the USA (Figure 1A), and another in the Middle East + Northeast Africa region (Figure 1B). The richness map for the future (Figure 2) shows that patterns of richness are predicted to change under future climatic conditions, where the main changes will be in the hotspot areas. The USA hotspot is predicted to reduce its area considerably due to the low species overlap in the future. By contrast, The Middle East + Northeast Africa hotspot is predicted to increase in size.

**FIGURE 2.**
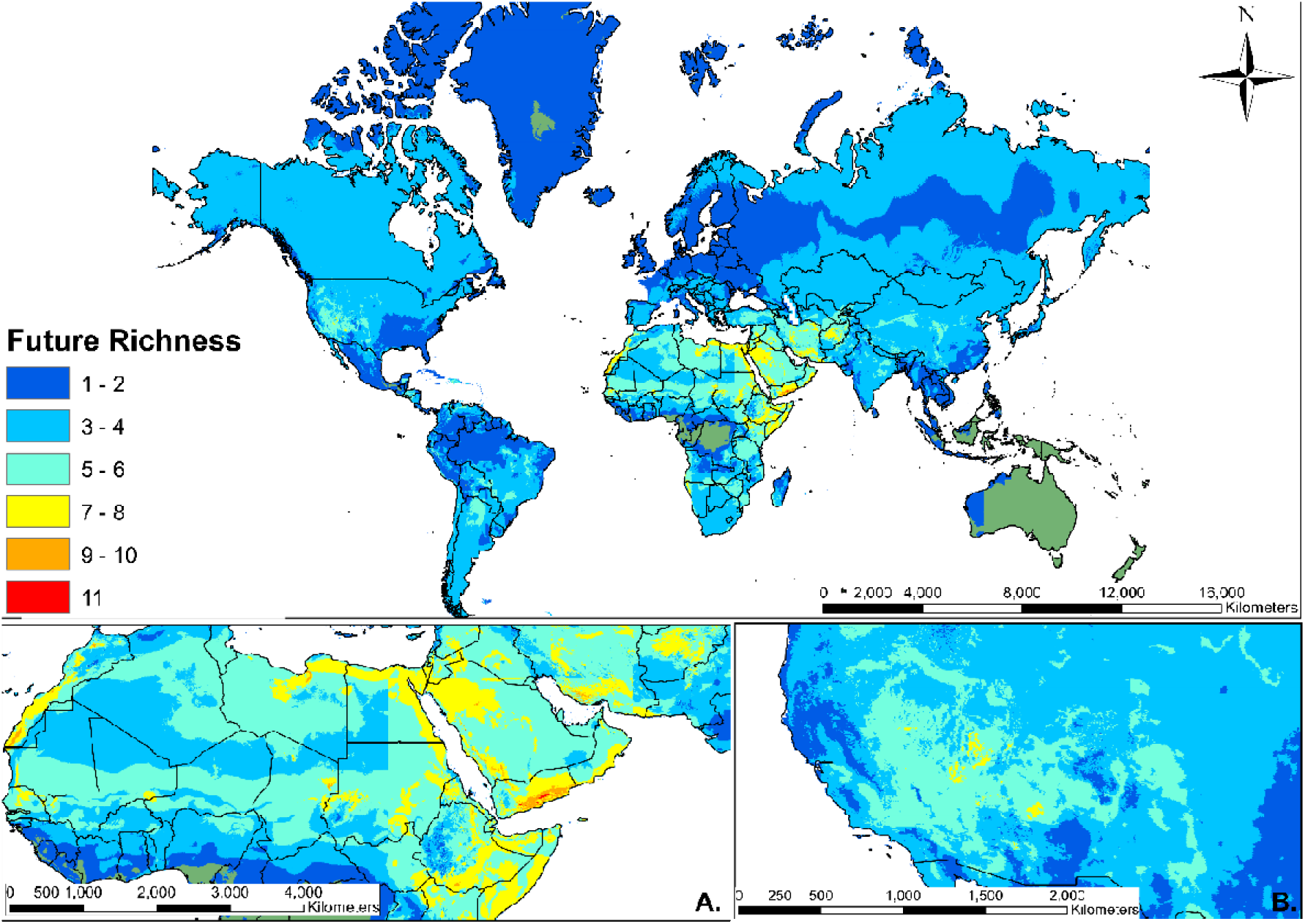
Species richness map of Canidae under future climate conditions produced by ENM. The richest areas (hotspots) were identified in the Middle East + Northeast Africa region (A) and western part of the USA (B). The legend on the left shows the number of overlapping species.

The ENMs indicated species that do not overlap currently will start to overlap in their distributions, and even those that overlap in only small parts of their distributions will suffer considerable increases in their overlap areas. In South America, *C. thous* is predicted to invade areas where only *A. microtis* and *Speothos venaticus* live, inside the Amazon rainforest (Appendix - Figures S39, S42, and S46). With the great expansion of *V. vulpes’* geographical distribution, this species is expected to overlap its area with *V. bengalensis*, *Vulpes rueppellii*, and *Vulpes zerda* (Appendix - Figures S45, S47, S48, and S49). In addition, *C. lupus* will probably overlap in areas occupied before only by *V. bengalensis*, *V. rueppellii*, *V. zerda*, and *Canis aureus* (Appendix - Figures S44, S47, S48, S49, and S50).

### 3.2 Evolutionary rescue

Most of canids presented evolutionary rates around 0.01 Haldanes (Table S2). The highest *H* value was found for *A. microtis* (*H* = 0.047 Haldanes), and the lowest value was from *Lycalopex griseus* (*H* = 0.004 Haldanes) (Table S2).

We found a significant weak negative correlation between change in range size and evolutionary potential: species that are predicted to undergo more habitat loss according to the ENMs have a lower potential for ER, according to the *H* values (Figure 3).

**FIGURE 3.**
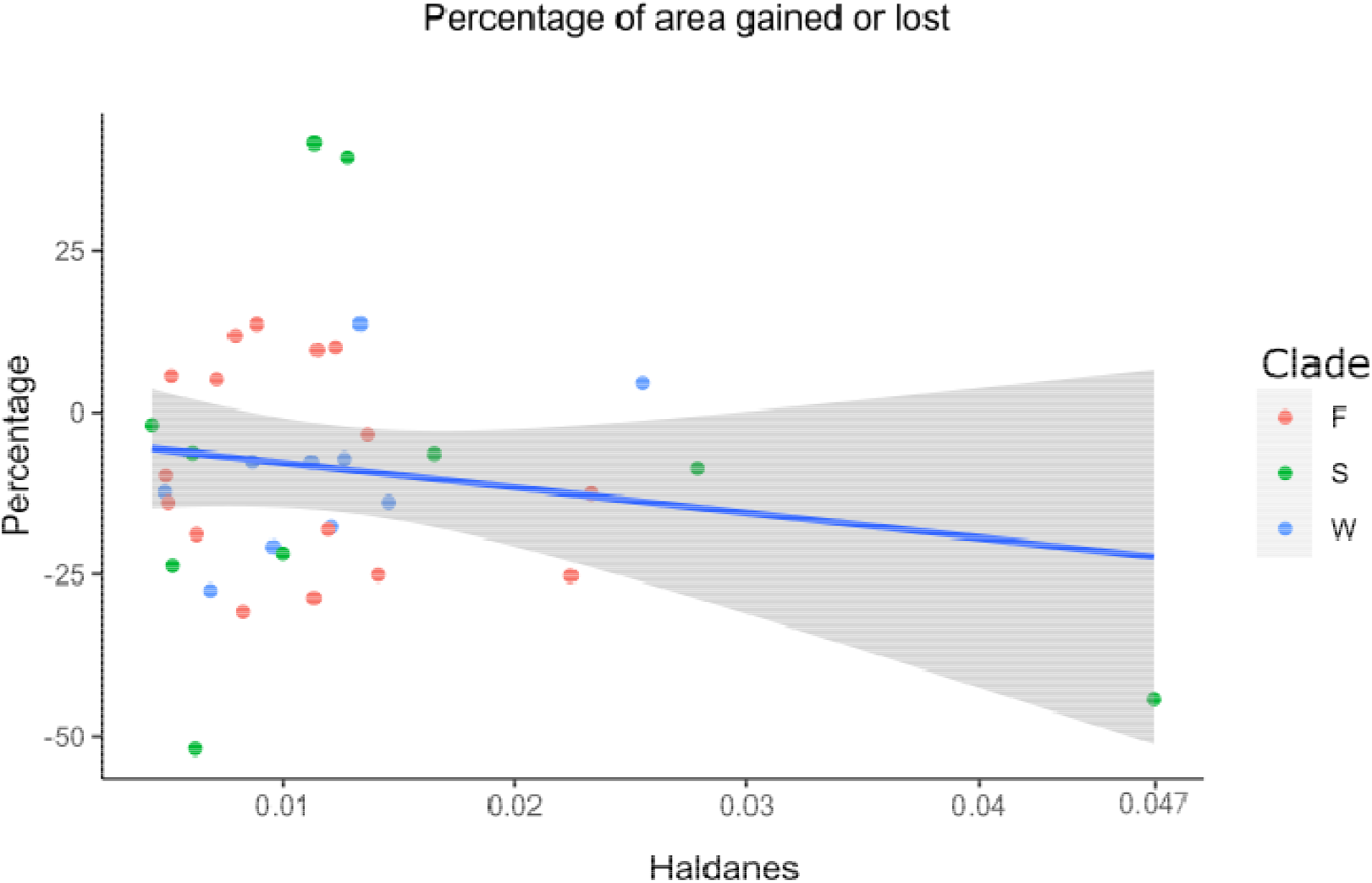
Plot representing the relationship between the percentage of area gained or lost by canids in relation to *H* values. The higher the *H* value, the lower the likelihood of evolutionary rescue. Red, green, and blue dots are species from the clades of foxes, South-American canids, and wolves, respectively. *R^2^* = −0.187 (*P* < 0.05).

## 4 DISCUSSION

We applied models of evolutionary rescue, using temperature and generation cycle as intrinsic characteristics of canids, together with ENMs to understand the magnitude of the effects of climate change on Canidae distribution. Predictions for the future by ENMs, derived from the IPCC worst climate change scenario, suggested that climate change will affect canids in distinct ways, where some species will expand or maintain their distributions, while most will suffer a large reduction in their suitable areas. Furthermore, the calculated Haldanes suggest that for some species it will be more difficult to keep up with the pace of temperature changes than others. We detected a weak negative correlation between habitat loss and potential for evolutionary rescue, indicating that the species with higher potential to evolutionary rescue are the ones that gain area or lose only a small part of their future distributions, while the ones which are going to lose a large part of their future distribution will need a higher evolutionary change to maintain their populations. *Atelocynus microtis*, for example, is predicted to lose about half of its potential distribution and has the highest *H* value among canids (*H* = 0.047 Haldanes). This negative correlation is to be expected because larger differences between present and future temperatures will increase *H* and will also make it more likely that range sizes will change.

Our results suggest that global warming will be devastating to the Canidae family as a whole. However, even in this pessimistic scenario, some species have the potential to benefit from future conditions and considerably expand their geographic distributions. In general, several taxa, including mammals, birds, amphibians, and reptiles, are expected to experience drastic range reductions (Araújo & New, 2007; Diniz-filho et al., 2009; Hidasi-neto et al., 2019; Lawler et al., 2009; Maiorano et al., 2011; Peterson et al., 2002). In a scenario such as this, several communities will probably lose phylogenetic and functional diversity (Davis et al., 1998; Hidasi-neto et al., 2019), and considering the number of interactions that will be lost within these areas, the ecological impacts due to indirect effects may be stronger than the direct effects of climate change on species’ distributions (Davis et al., 1998; Peterson et al., 2002). Carnivores, through population regulation, can promote the coexistence of several species by reducing interspecific competition (Paine, 1966). Because canids, being carnivores, hunt distinct animals, they end up regulating the population dynamics of their prey, which is an important factor for maintaining biodiversity (Sanders et al., 2013; Sanders & van Veen, 2012).

In South-America, there is a very concerning situation, where *A. microtis* will probably contract its range substantially and undergo fragmentation of its distribution within the Amazon Forest, while *C. thous* will expand. *A. microtis* is ecologically restricted to very specific resources and conditions (Sillero-Zubiri et al., 2004; Wilson & Mittermeier, 2009). By contrast, *C. thous* is a generalist species with a large distribution across South-America (Sillero-Zubiri et al., 2004). Currently, the status of *A. microtis* is “Near Threatened” (IUCN, 2019), but considering the climate change effects shown here, and the fact that the Amazon Forest has been suffering with wildfires and an intense deforestation process over the last decades (Exbrayat et al., 2017; INPE, 2019), *A. microtis* is probably experiencing a substantial habitat loss followed by a very likely increase in the number of direct encounters with another competitor. Thus, we suggest that its “Near Threatened” status must change, at least, to “Vulnerable”.

A similar situation applies to *V. vulpes* and *V. lagopus*. The first one has a wide distribution over the northern hemisphere, while the second is restricted to areas covered by snow around The North Pole, but both species overlap in the Tundra of North America and Eurasia (Hersteinsson & Macdonald, 1992; Sillero-Zubiri et al., 2004). Over the past few years there has been an increase in the number of encounters between the two species due to the warming temperatures that are gradually melting the Arctic ice cap, reducing the available area for *V. lagopus*, but making it possible for *V. vulpes* to expand its distribution to the north into arctic tundra in Eurasia and North America (Gallant et al., 2012). This reality is even more aggravating in the future scenario shown here, considering the large area loss by *V. lagopus* to *V. vulpes* (Figure S51). However, Gallant et al. (2012) suggested that food scarcity in these areas seems to explain the dynamics of the geographical overlap of both two species better than climate warming. Nevertheless, the effects of area loss must still be taken into account to outline conservation strategies for *V. lagopus*.

The loss of species has severe impacts on the functioning of ecosystems (Cardinale et al., 2012; Kennedy et al., 2002; Lyons & Schwartz, 2001; Pimm et al., 2014). In general, reductions in the number of species (functional groups) decrease the efficiency of communities to capture resources, and convert these into biomass (Balvanera et al., 2006; Cardinale et al., 2012; Quijas et al., 2010). Our niche models detected two major richness hotspots for Canidae: one in the Middle East + North East Africa and one in North America. The former is predicted to undergo a small expansion, mainly due to the range expansion of *C. lupus*, *C. anthus*, and *V. vulpes* over these areas, and the capacity of *C. lupus* and *V. vulpes* to live around urban areas (Sillero-Zubiri et al., 2004; Wang & Tedford, 2008; Wilson & Mittermeier, 2009). This capacity can also explain the wide distribution of both species around the world. The other hotspot area, in North America, is expected to experience a considerable area reduction. This can be explained by the small portion of this hotspot that is within protected areas in the USA, according to Brum et al. (2017).

Here, the ENMs for all canids (appendix) agreed well with the current distribution of canids, suggesting that the methodology we applied is reliable to assess the impacts of climate change on Canidae, taking into account their main niche dimensions. *Canis lupus* is the only species for which the ENMs for the present did not encompass the entire distribution presented by its polygon, because it is not predicted to occur in the Middle East. This might be explained by the presence of a single population found in that region, which results in the distribution of the species to be extended to areas that are not suitable. The IUCN distribution maps are widely used in several studies for different purposes (Kyne et al., 2020; Porto et al., 2021; Shier, 2015; Zhang et al., 2019), and are defined as the area within the outermost limits of known occurrence for a species, but this area is not an estimate of the extent of occupied habitat, it only measures the general extension of the localities in which the species is found (Gaston & Fuller, 2009). Thus, polygons are highly susceptible to sampling biases. Nevertheless, it is important to point out that ENMs for the future suggest that *Canis lupus* will expand its distribution to the Middle East, which could be an indication that this region is already becoming suitable for the species.

Our methods assumed that the prey of the Canidae will respond to environmental changes at the same rate as their (apex or medium-level) predators. Indeed, climate change has already been observed to have wide-ranging trophic effects (Gilman et al., 2010), and physiological and behavioral effects in other species (Parmesan, 2006). Modelling the effect of climate change on species’ communities and trophic interactions has proven extremely difficult, but these interactions can have serious impacts on species distributions (Sanders et al., 2013; Sanders & van Veen, 2012). These trophic interactions may be further disrupted by invasive species, the spread of which could be accelerated by climate change (Hellmann et al., 2008).

Looking at the *H* values, two cases are very concerning. *Atelocynus microtis* and *Chrysocyon brachyurus* present higher *H* values compared to other canids (0.047 and 0.027, respectively), and based on Diniz-Filho et al. (2019), these species have a lower potential for evolutionary rescue. Although *H* values and ENMs try to elucidate the future of species, they have distinct points of view about the effects of climate change on canids, and therefore should not be compared. However, these two approaches can shed light on Canidae responses to the future of the planet. *H* values suggest that some species have less potential than others to adapt fast enough to temperature changes, but ENMs indicate that some of them may increase their range, because more suitable habitats will become available for them due to climate change. Thus, in these cases ecological processes seem to prevail over evolutionary ones.

Unfortunately, very little is known about ER in nature to compare with our findings, mostly because the idea that evolution may influence the persistence of a population facing a rapid environmental change is very recent. Nevertheless, Diniz-Filho et al. (2019) already suggested that the use of the ER approach for wider geographical areas might not be that simple. They suggested that in order to obtain a standard temperature deviation, the real temperature tolerances must be known. However, no lab values were available for any wild canid, meaning that only values obtained from range estimations and ENMs could be used. Nonetheless, both may underestimate a species’ true temperature tolerance. For example, while we have extracted values of mean Bio05 (maximum temperature in the warmest month), sometimes these values are well below the highest value seen within a species range.

The biogeographic patterns observed in this study may provide useful information for assessing how canids are distributed in the present over the planet, being an alternative to the distribution polygons provided by IUCN (2020). Climate change is projected to play an essential role in the geographical distribution of canids, so our predictions can be used to identify key areas for conservation strategies. This should receive special attention because as we showed, most of the Canidae hotspot regions are not located within protected areas.

## DECLARATION SECTION

### Ethics approval and consent to participate

Not applicable.

### Consent for publication

Not applicable.

### Availability of data and material

All data generated or analyzed during this study are included in this article and in the supplementary files.

### Authors’ contributions

L.M.V.P. and D.B. conceived and designed the study and analyses. L.M.V.P. and D.B. performed the analyses. L.M.V.P. and D. B. wrote the first draft of the manuscript. R.S.E. and R.M. commented on the methods and contributed to substantial revisions on the draft.

## SUPPLEMENTARY TEXT: FIGURE LIST

**FIGURE S38.**
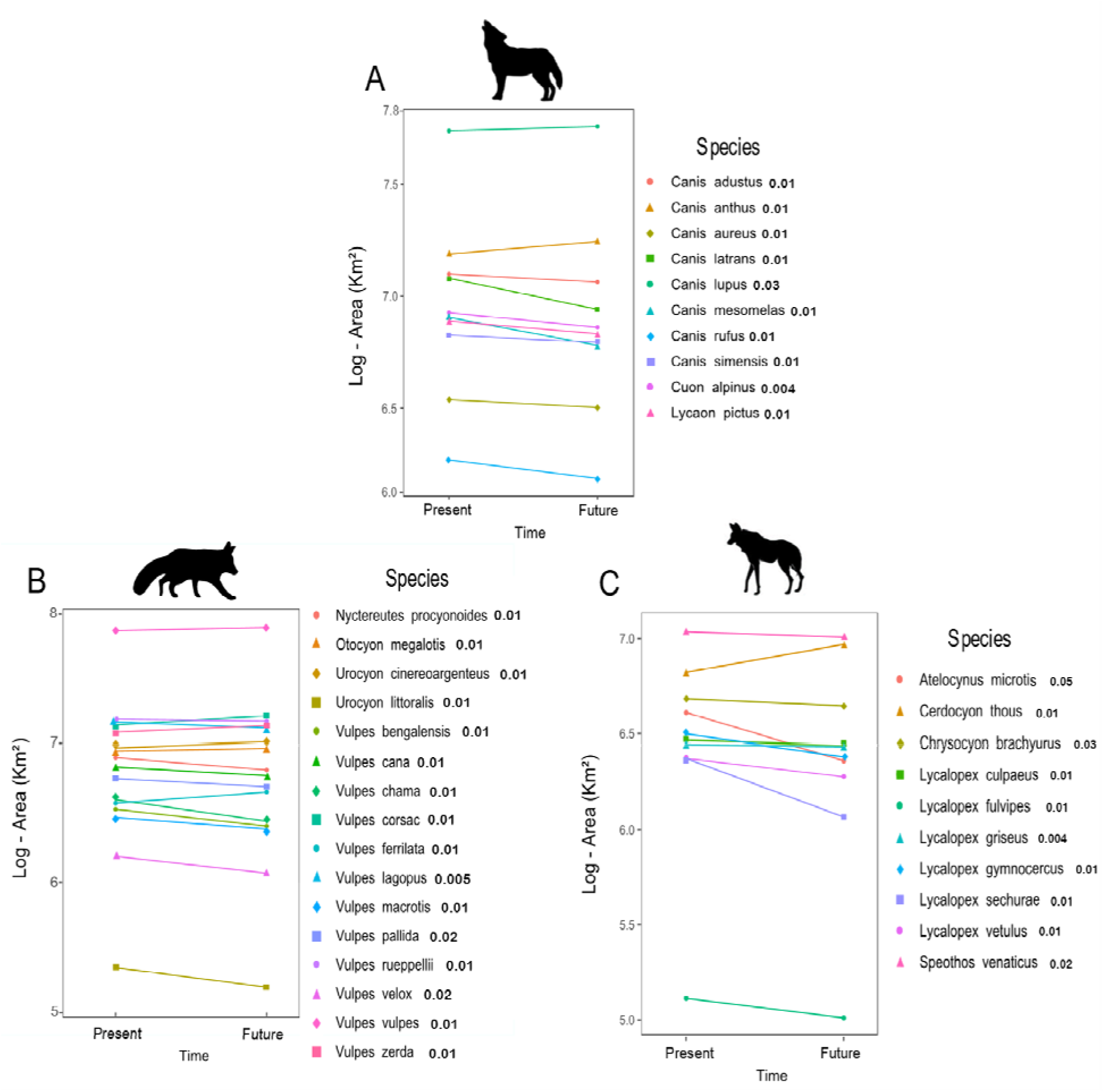
Plot representation, on logarithmic scale, of range expansion or contraction over time for the clades of wolves (A), foxes (B), and South American canids (C). *H* values for each species are indicated next to each species name.

**FIGURE S51.**
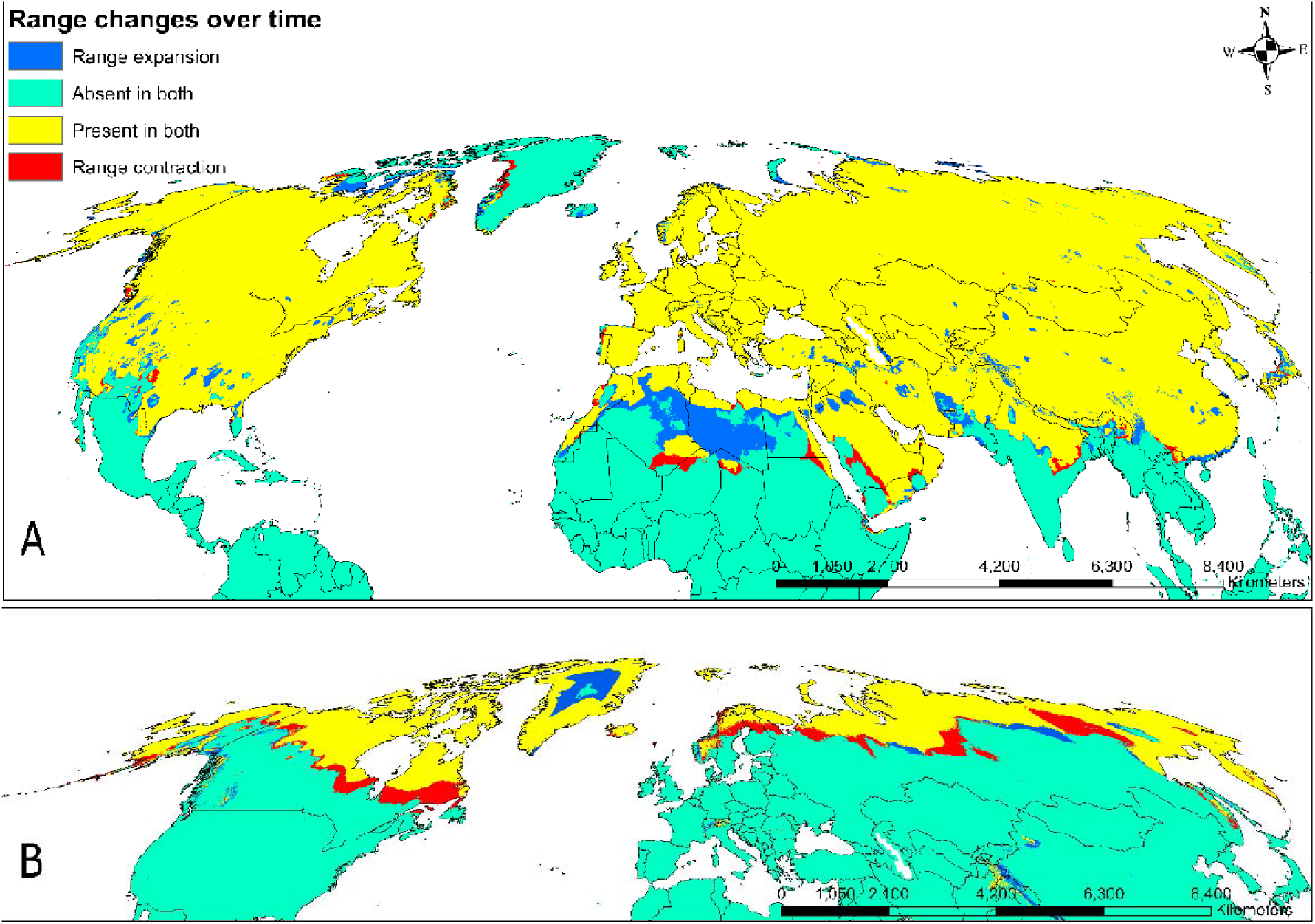
Comparison of present and future suitable areas of *Vulpes vulpes* (A) and *Vulpes lagopus* (B). The image shows regions where loss is expected to occur (red) and regions where the species will increase their distributions (blue).

## SUPPLEMENTARY TEXT: TABLE LIST

**TABLE S1.** List of the 36 species of Canidae included in our study. Age of sexual maturity of females (years), the number of generations until 2075, the number of occurrence points for each species, and the source of the original description are indicated here.

| Species | Sexual maturity of females | Number of generations | Number of occurrence points | Descriptor |
| --- | --- | --- | --- | --- |
| *Canis adustus* | 0.75 | 100 | 1.028 | Sundevall, 1847 |
| *Canis aureus* | 1 | 75 | 2.769 | Linnaeus, 1758 |
| *Canis anthus* | 1 | 75 | 1.536 | Cuvier, 1820 |
| *Canis lupus* | 2.5 | 25 | 8.490 | Linnaeus, 1758 |
| *Canis latrans* | 0.84 | 89.3 | 2.402 | Say, 1823 |
| *Canis mesomelas* | 0.84 | 89.3 | 645 | Schreber, 1775 |
| *Canis rufus* | 1 | 75 | 30 | Audubon & Bachman, 1851 |
| *Canis simensis* | 2 | 37.5 | 12 | Rüppell, 1840 |
| *Cuon alpinus* | 1 | 75 | 507 | Pallas, 1811 |
| *Lycaon pictus* | 1.23 | 60.4 | 281 | Temminck, 1820 |
| *Nyctereutes procyonoides* | 0.82 | 91.5 | 846 | Gray, 1834 |
| *Vulpes bengalensis* | 1.5 | 50 | 327 | Shaw, 1800 |
| *Vulpes cana* | 0.82 | 91.5 | 396 | Blanford, 1877 |
| *Vulpes chama* | 0.75 | 100 | 229 | A. Smith, 1833 |
| *Vulpes corsac* | 1.38 | 54.3 | 1.193 | Linnaeus, 1768 |
| *Vulpes ferrilata* | 1.15 | 65.2 | 264 | Hodgson, 1842 |
| *Vulpes macrotis* | 0.82 | 91.5 | 229 | Merriam, 1888 |
| *Vulpes pallida* | 1 | 75 | 406 | Cretzschmar, 1826 |
| *Vulpes rueppellii* | 1 | 75 | 1.299 | Schinz, 1825 |
| *Vulpes velox* | 1 | 75 | 88 | Say, 1823 |
| *Vulpes vulpes* | 0.83 | 90.4 | 9.457 | Linnaeus, 1758 |
| *Vulpes zerda* | 0.49 | 153.1 | 850 | Zimmermann, 1780 |
| *Vulpes lagopus* | 0.83 | 90.4 | 3.468 | Linnaeus, 1758 |
| *Urocyon cinereoargenteus* | 0.95 | 78.7 | 1.089 | Schreber, 1775 |
| *Urocyon littoralis* | 1 | 75 | 30 | Baird, 1857 |
| *Otocyon megalotis* | 0.61 | 122.6 | 515 | Desmarest, 1822 |
| *Atelocynus microtis* | 1 | 75 | 238 | Sclater, 1883 |
| *Cerdocyon thous* | 0.76 | 98.7 | 864 | Linnaeus, 1766 |
| *Chrysocyon brachyurus* | 2 | 37.5 | 457 | Illiger, 1815 |
| *Lycalopex culpaeus* | 1 | 75 | 345 | Molina, 1782 |
| *Lycalopex fulvipes* | 1 | 75 | 8 | Martin, 1837 |
| *Lycalopex griseus* | 1 | 75 | 255 | Gray, 1837 |
| *Lycalopex gymnocercus* | 1 | 75 | 312 | G. Fischer, 1814 |
| *Lycalopex sechurae* | 1 | 75 | 24 | Thomas, 1900 |
| *Lycalopex vetulus* | 1 | 75 | 183 | Lund, 1842 |
| *Speothos venaticus* | 0.83 | 90.4 | 1.076 | Lund, 1842 |

**TABLE S2.** Area difference in species distributions for present and future, showing expansion or retraction of canids’ geographical distributions. *H* values are also indicated.

| Species | Present area (Km<sup>2</sup>) | Future area (Km<sup>2</sup>) | *H* |
| --- | --- | --- | --- |
| *Atelocynus microtis* | 4.379.627 | 2.438.970 | 0,047520 |
| *Canis anthus* | 15.472.384 | 17.583.018 | 0,013342 |
| *Canis aureus* | 3.448.559 | 3.187.594 | 0,008698 |
| *Canis latrans* | 12.065.866 | 8.749.988 | 0,006891 |
| *Canis lupus* | 55.058.863 | 57.577.546 | 0,02551 |
| *Canis mesomelas* | 7.840.423 | 6.214.900 | 0,009592 |
| *Canis rufus* | 1.858.413 | 1.529.932 | 0,012079 |
| *Canis simensis* | 6.707.343 | 6.227.124 | 0,012671 |
| *Canis adustus* | 12.577.073 | 11.623.101 | 0,011225 |
| *Cerdocyon thous* | 7.224.726 | 10.225.538 | 0,011357 |
| *Chrysocyon brachyurus* | 5.202.737 | 4.755.462 | 0,027895 |
| *Cuon alpinus* | 7.757.856 | 6.803.830 | 0,004926 |
| *Lycalopex culpaeus* | 3.121.231 | 2.923.795 | 0,006086 |
| *Lycalopex fulvipes* | 126.236 | 98.762 | 0,009984 |
| *Lycalopex vetulus* | 2.539.881 | 2.040.550 | 0,012808 |
| *Lycalopex griseus* | 2.961.540 | 2.903.216 | 0,004344 |
| *Lycalopex gymnocercus* | 3.354.884 | 2.561.320 | 0,005269 |
| *Lycalopex sechurae* | 2.514.432 | 1.209.669 | 0,006207 |
| *Lycaon pictus* | 8.445.869 | 7.276.908 | 0,014579 |
| *Nyctereutes procyonoides* | 7.413.459 | 6.018.161 | 0,006262 |
| *Otocyon megalotis* | 8.251.366 | 8.676.914 | 0,007170 |
| *Speothos venaticus* | 11.953.879 | 11.185.765 | 0,016549 |
| *Urocyon cinereoargenteus* | 8.757.468 | 9.595.434 | 0,011490 |
| *Urocyon littoralis* | 200.615 | 143.194 | 0,011346 |
| *Vulpes bengalensis* | 3.053.463 | 2.287.423 | 0,014108 |
| *Vulpes cana* | 6.315.447 | 5.439.834 | 0,005072 |
| *Vulpes chama* | 3.594.029 | 2.487.370 | 0,008272 |
| *Vulpes corsac* | 13.114.501 | 14.423.740 | 0,012275 |
| *Vulpes ferrilata* | 3.502.426 | 3.977.712 | 0,008895 |
| *Vulpes lagopus* | 13.405.437 | 12.101.093 | 0,004969 |
| *Vulpes macrotis* | 2.651.764 | 2.171.680 | 0,011933 |
| *Vulpes pallida* | 5.164.447 | 4.518.576 | 0,023281 |
| *Vulpes velox* | 1.360.294 | 1.016.829 | 0,022411 |
| *Vulpes vulpes* | 64.415.599 | 68.080.936 | 0,005214 |
| *Vulpes zerda* | 11.242.325 | 12.574.885 | 0,007949 |
| *Vulpes rueppellii* | 14.074.266 | 13.588.853 | 0,013633 |

